# Engineering neurovascular organoids with 3D printed microfluidic chips

**DOI:** 10.1101/2021.01.09.425975

**Authors:** Idris Salmon, Sergei Grebenyuk, Abdel Rahman Abdel Fattah, Gregorius Rustandi, Thomas Pilkington, Catherine Verfaillie, Adrian Ranga

**Affiliations:** Laboratory of Bioengineering and Morphogenesis, Biomechanics Section, Department of Mechanical Engineering, KU Leuven, Leuven, Belgium; FabLab Leuven, KU Leuven Research & Development Belgium; Stem Cell and Developmental Biology, Department of Development and Regeneration, KU Leuven, Leuven, Belgium

## Abstract

The generation of tissues and organs requires close interaction with vasculature from the earliest moments of embryonic development. Tissue-specific organoids derived from pluripotent stem cells allow for the in vitro recapitulation of elements of embryonic development, however they are not intrinsically vascularized, which poses a major challenge for their sustained growth and for understanding the role of vasculature in fate specification and morphogenesis. Current organoid vascularization strategies do not recapitulate the temporal synchronization and spatial orientation needed to ensure in-vivo-like early co-development. Here, we developed a human pluripotent stem cell (hPSC)-based approach to generate organoids which interact with vascular cells in a spatially determined manner. The spatial interaction between organoid and vasculature is enabled by the use of a custom designed 3D printed microfluidic chip which allows for a sequential and developmentally matched co-culture system. We show that on-chip hPSC-derived pericytes and endothelial cells sprout and self-assemble into organized vascular networks, and use cerebral organoids as a model system to explore interactions with this de novo generated vasculature. Upon co-development, vascular cells interact with the cerebral organoid and form an integrated neurovascular organoid on chip. This 3D printing-based platform is designed to be compatible with any organoid system and is an easy and highly cost-effective way to vascularize organoids. The use of this platform, readily performed in any lab, could open new avenues for understanding and manipulating the co-development of tissue-specific organoids with vasculature.

## Introduction

Organoids derived from both adult and pluripotent stem cells have emerged in recent years as in-vitro model systems which can recapitulate key features of physiology, with applications in drug screening and regenerative medicine (1). In particular, organoids derived from human pluripotent stem cells (hPSC) have also recapitulated key features of human embryonic development and thereby provided an unprecedented view into early human development (1,2). However they have been limited in size, reproducibility and complexity.

The presence of a vascular network has been hypothesized to be a key missing feature in these models which could account for some of these limitations. Indeed, the endothelium plays an critical role in the development of tissues and organs, functioning as a site of nutrient and waste exchange as well as a paracrine signalling centre which directs differentiation, patterning and morphogenesis (3). For example, the endothelium plays an in important role in the initiation of cortical neurogenesis as well as in the outgrowth of the liver bud by surrounding the embryonic hepatic endoderm, thereby delimitating the mesenchymal domain (4). In the immature brain, a relief of hypoxia mediated by endothelium invasion is shown to trigger neural stem cell differentiation (5). As such, this endothelial cell (EC) co-development would be a critical and necessary component for the biomimetic development of in-vitro tissues such as organoids.

Early attempts at organoid vascularization focused on the transplantation of these organoids into murine hosts (6,7). Intestinal as well as cerebral organoids were shown to integrate and mature well into the host, with host vasculature seen to invade the organoid followed by the establishment of a functional perfused vascular network. While these observations underscore the importance of vasculature in the development of human organoids, the use of an in-vivo host with endogenous vasculature is clearly not a practical approach for organoid vascularization, prompting a number of attempts at in-vitro vascularization. The most common approach has involved the co-culture of organoids, including liver, pancreas, colon, kidney and cerebral organoids with primary ECs, with human umbilical cord endothelial cells (HUVEC) being most frequently used (8–12). However HUVECs are already developmentally specified in contrast to hPSC-derived ECs, which can interact and specify the maturation of specific tissues. Indeed, liver 2D and 3D vascular co-culture systems harbouring hPSC-derived ECs showed higher expression of CYP enzymes and albumin when compared to co-cultures containing HUVECs (13,14).

Recent advances in organoid vascularization have also focussed on the use of hPSCs engineered to overexpress the EC-specific ETV2 transcription factor. Co-aggregation of engineered and wild-type cells followed by neural differentiation with overexpression of ETV2 led to the generation of cerebral organoids with interspersed self-assembled vascular cells and enhanced functional maturation, including rudimentary blood brain barrier characteristics (15). Such co-culture models where vascular and cerebral cells are continuously grown together do not, however, mimic the characteristic spatial organization and temporal synchronization of angiogenic embryonic development, where blood vessels invade into an initially avascular neuroectodermal tissue (16–18). Microfluidic devices characterized by fluidically linked microchannels are ideally suited to provide such spatial interactions. In these devices, cells which are initially cultured separately can interact in designed spaces by migration through hydrogel substrates within the channels (19,20). However, an important limitation to the broader applicability of such devices for biological applications is that the soft lithography technologies necessary to fabricate these devices require access to dedicated clean room equipment and are time-consuming and error-prone, leading to a slow process for design iteration and optimization. High resolution 3D printing technology can overcome these limitations by offering the ability to control the spatial environment in a versatile, on-demand and easy to use manner (21,22). In particular, 3D printing technologies based on stereolithography (SLA) have seen in recent years a remarkable increases in spatial resolution and in availability of new printable resin formulations, including those which can be rendered biocompatible with post-printing processing (23,24). These technologies are now readily accessible to most labs due to significant price drops in both hardware and materials in recent years (25). Here, we explore the use of consumer-grade 3D printing technologies as a key enabling technology to generate temporally and spatially synchronized all hPSC vascularized organoid.

### 3D printing for microfluidic organoid vascularization

In order to tackle the challenge of both temporal and spatial synchronization between organoid and vasculature, we developed an approach based on two key aspects: the use of all-hPSC derived cell lineages and the development of customized 3D printing to engineer microfluidic chips with designed geometries. We hypothesized that by simultaneously initiating vascular and neural differentiation from PSCs, we would obtain developmentally matched lineages, which could then be grown in a spatially defined manner on a biomimetic chip (**Fig 1A**).

**Figure 1.**
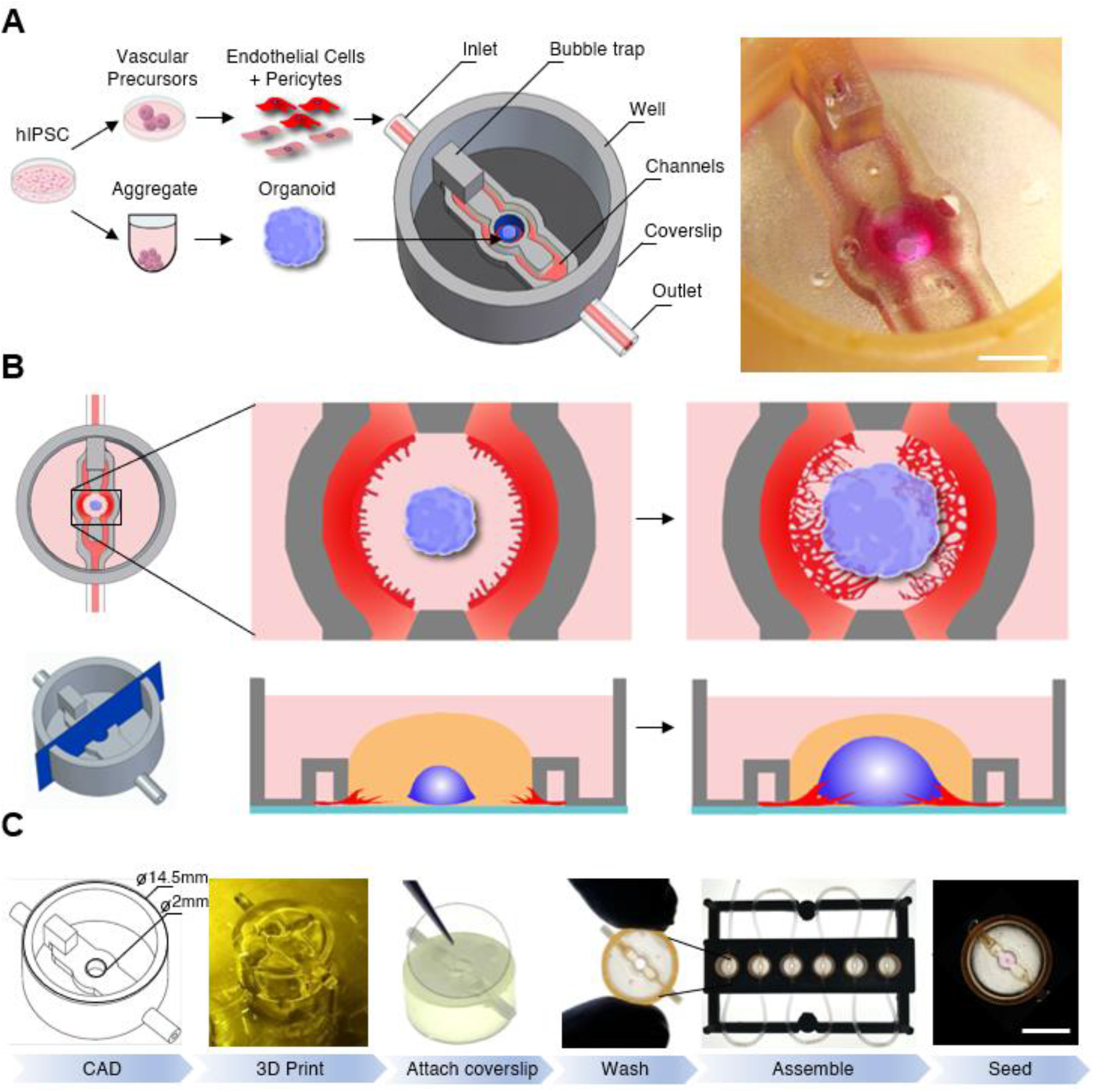
3D printed microfluidic platform for vascularized organoid cultures on chip. (**A**) Schematic flow for initiation of interaction culture involving differentiation of hPSC into vascular cells and early neural organoids in suspension followed by seeding into 3D printed microfluidic chip, and stereomicroscope image of the organoid on chip (scale bar: 2mm). (**B**) Schematic representations of on-chip angiogenic sprouting resulting in a vascularized organoid on chip. (**C**) Microfluidic chip manufacturing process for multiplexed mono and vascularized co-cultures on chip (CAD: Computer-aided design), scale bar: A: 2mm, C: 5mm.

A key feature of our microfluidic chip is its “open well” design. In contrast to commonly used microfluidics where the culture chamber is closed, this design does not have any physical seal over the main culture chamber, thereby allowing for easy and direct accessibility to this compartment (12,26–28). In the context of organoid culture, this design therefore allows for direct and precise placement of an organoid in the central compartment as well as its easy removal for downstream analysis such as immunostaining. Another advantage of this design is that an individual organoid can be seeded in each chip, making it an ideal platform to study co-developmental processes at single-organoid level (29).

To support vascular cell sprouting, network formation, and vascular invasion of a growing organoid on-chip, the open well microfluidic chip is designed such that microfluidic channels flanking the central organoid chamber can be seeded with vascular cells via an inlet (**Fig. 1A,1B**). Communication of diffusible molecules and cells between the organoid chamber compartment and the microfluidic channels is ensured by a nearly complete wall separating the channels and the main compartment, leaving only a 50μm gap between the coverslip and the 3D printed microfluidic chip (**Fig.1B bottom**).

In order to print our parts, we sought an enabling printing technology and associated material which would allow both high-resolution printing (circa 200μm feature size) as well as biocompatibility. **(Supplementary Fig. 1A)**. We choose the Formlabs Form 2 printer, a consumer-grade 3D instrument based on stereolithography technology equipped with a 140 μm spot size laser which allows for printing in the micrometer range and can print a variety of biocompatible printing materials. In particular, the Dental SG material, which has been reported to show no toxicity when exposed to 3T3 and hepatoma cells, was selected as the material to print our microfluidic devices (24). To render the chip suitable for microscopy, we designed and printed a bottomless chip (**Fig. 1A)**, which was completed in a subsequent step by the permanent attachment of glass coverslip to its bottom using a biocompatible UV-curable glue (**Fig. 1C)**.

Our initial experiments indicated that hPSC culture was not compatible with the printed chips, suggesting remaining toxicity of the printed material. To render the chip biocompatible, we therefore developed an extensive washing procedure which allowed the unreacted polymers to leach out. Taking advantage of the design flexibility of 3D printing technologies, we also optimized a chip fabrication pipeline that allowed us to process tens to hundreds of chips in the same day (**Fig.1C**). To parallelize the handling and image acquisition of experiments within these chips, we used inexpensive, lower resolution fused deposition modelling printing of Polylactic acid (PLA) plastic to fabricate plates capable of holding 6 microfluidic devices. This multiplexed platform was instrumental in allowing on-chip biological experiments in a standardized and reproducible manner.

### Vascular networks on 3D printed microfluidic chip

In order to produce cells for the vascular component of the chip, we aimed at generating both ECs and pericytes. Indeed, mural cell types such as pericytes play a crucial role in blood vessel stabilization *in vivo* and the inclusion of such cells in a microfluidic vascularization chip would therefore present an important advantage for long term vascular culture on chip (30). A recently published protocol demonstrated the directed differentiation of hPSCs towards dual EC and pericyte fates in the form of 3D blood vessel organoids (31). We adapted this protocol to an initially two-dimensional differentiation in order to rapidly obtain a large cell yield for microfluidic chip seeding (**Fig.2A; Supplementary Fig. 2A**). In order to confirm that this 2D differentiation protocol yields both ECs and pericytes, immunohistochemistry (IHC) was performed at day 6 of differentiation, prior to cell dissociation and immediate seeding into the microfluidic chip. The presence of both pericytes, marked by PDGFRβ and ECs, marked by CD31 was confirmed at this time, thereby ensuring that the cells collected for seeding into the chip contained both vascular lineages (**Fig.2A**). This mixed vascular cell population was then used for seeding the channels of the microfluidic chip. To provide a substrate for 3D vascular invasion, we filled the central organoid chamber with Matrigel, an extracellular matrix frequently used in angiogenic sprouting assays (32). Vascular cell sprouting was triggered by the addition of 100ng/mL of vascular endothelial growth factor A (VEGF-A) to the culture medium in the central compartment. Sprouting of both ECs and pericytes on-chip was observed by IHC at day 7 (**Fig.2B**). Vascular cells which were initially seeded in the microfluidic channels were observed to start sprouting into the organoid culture chamber via an intermediate zone (**Fig.2B**). These vascular cells were observed in the organoid culture chamber by day 10 of differentiation and progressively sprouted towards the center of the organoid culture chamber (**Fig.2C**). The length of these vascular sprouts was measured over 6 days, starting from the day when they first appeared in the organoid culture chamber, and was seen to increase over this observed period (**Fig.2C**). This invasion speed was constant over the observed period and was calculated to be on average 80 μm/day. To determine whether pericytes and ECs would maintain their identity over a longer differentiation period, we stained for PDGFβ and CD31 at day 15 and day 30. Clear differences in vascular organization were evidenced between day 15 and day 30 cultures (**Fig.2D-E)**. At day 15 the pericytes and EC had a mixed and disorganized appearance, with pericytes and ECs in the same plane (**Fig.2F top panel**). In contrast, at day 30, the vascular network was much further developed, with CD31+ cells contributing to consolidated vessel-like structures, and pericytes no longer evidenced in the same plane as the ECs and instead forming a supportive layer above the formed vascular networks (**Fig.2G; Supplementary Fig.2B**).

**Figure 2.**
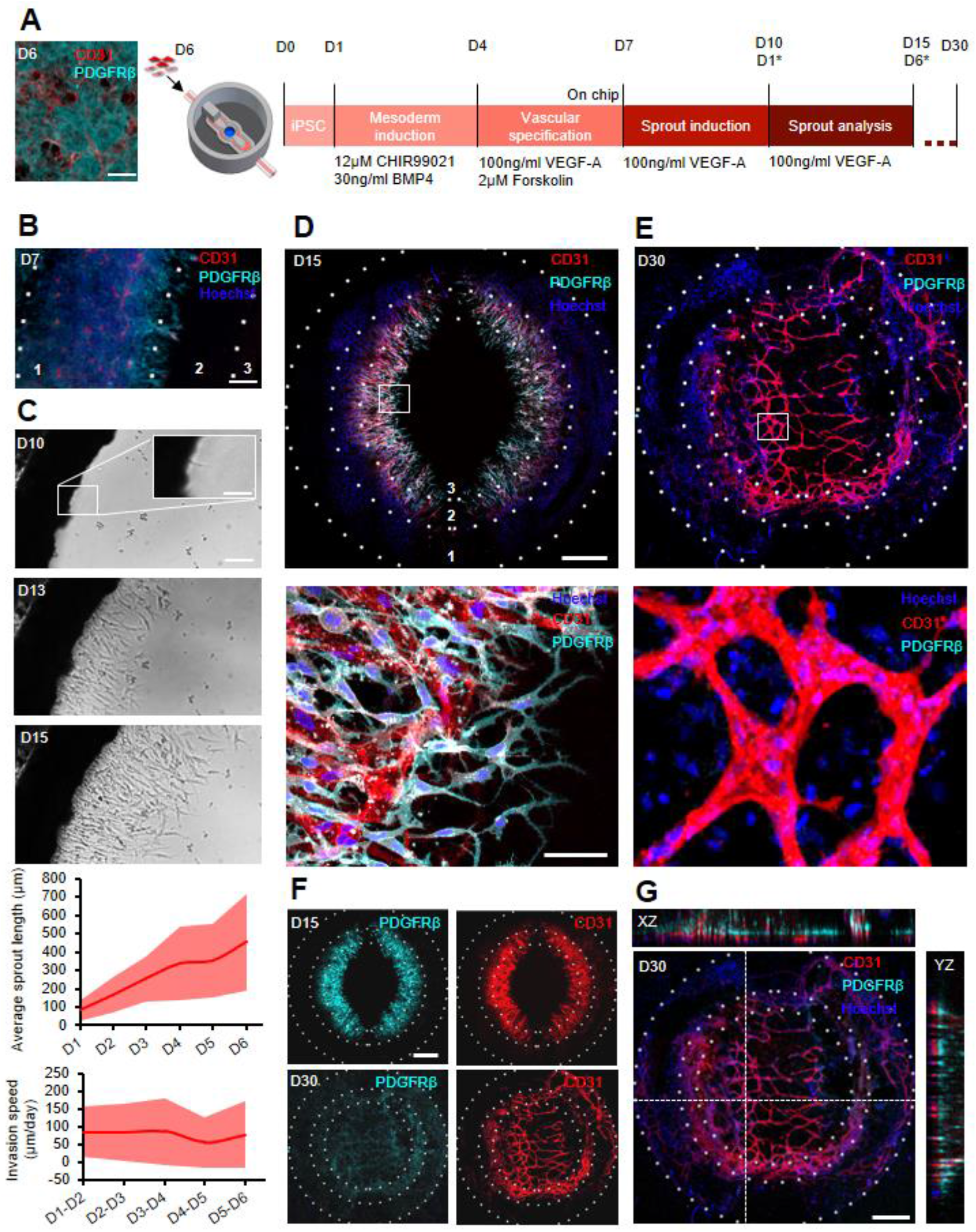
Vascular networks on 3D printed microfluidic chip. **(A)** Differentiation protocol for the generation of hPSC derived vascular networks on chip. (**B**) IHC image of vascular cells on chip at day 7 with sprouts present in the interface (2) between channels (1) and the organoid culture chamber (3) (scale: 100μm). (**C**) Bright field images of the organoid culture chamber showing sprout progression and graphs quantifying sprout progression (n=3;Avg.;SD) with 4-6 chips analysed per n (scale: 100μm, 40μm for inset) **(D**) IHC of D15 vascular culture on chip showing presence of CD31+ ECs and PDGFRβ+ pericytes (scale: 500μm). (**E**) IHC of D30 vascular networks on chip (scale: 500μm). (**F**) IHC stains for PDGRRβ and CD31 at day 15 and day 30 (scale: 500μm). (**G**) Bottom slice of a 200μm stack of D30 vascular networks on chip with corresponding ZX an ZY confocal planes showing layered appearance of CD31 and PDGFRβ cells in two layers (scale: 500μm)

### On chip sprout progression and cerebral organoid development

In order to demonstrate that our vascularization strategy could be applied to organoid models, we chose to incorporate the widely used cerebral organoid model into our chips (33). The generated aggregates exhibited a circular translucent band at the edges after two days of neural differentiation (**Supplementary Fig.3A**), suggesting successful induction of hPSCs into neural epithelial cells (33). At day 5 of neural differentiation, these neural aggregates were embedded in Matrigel and seeded into the organoid chamber of the chip for further maturation and growth. Organoid growth on chip was quantified from bright field images acquired every two days, and was seen to increase over the entire course of differentiations (**Fig.3A**). Cerebral organoid maturation was verified by staining for Pax6, the earliest marker of human neural identity, at day 15 and βiii tubulin, a post mitotic neuron marker, at day 30 **(Supplementary Fig.3B**) (34). Organoids also exhibited characteristically complex internal cytoarchitecture consisting of multiple epithelial domains. Together, these results demonstrate that 3D printed chips support the growth, differentiation and morphogenesis of cerebral organoids, as well as the development of complex vascular networks.

**Figure 3.**
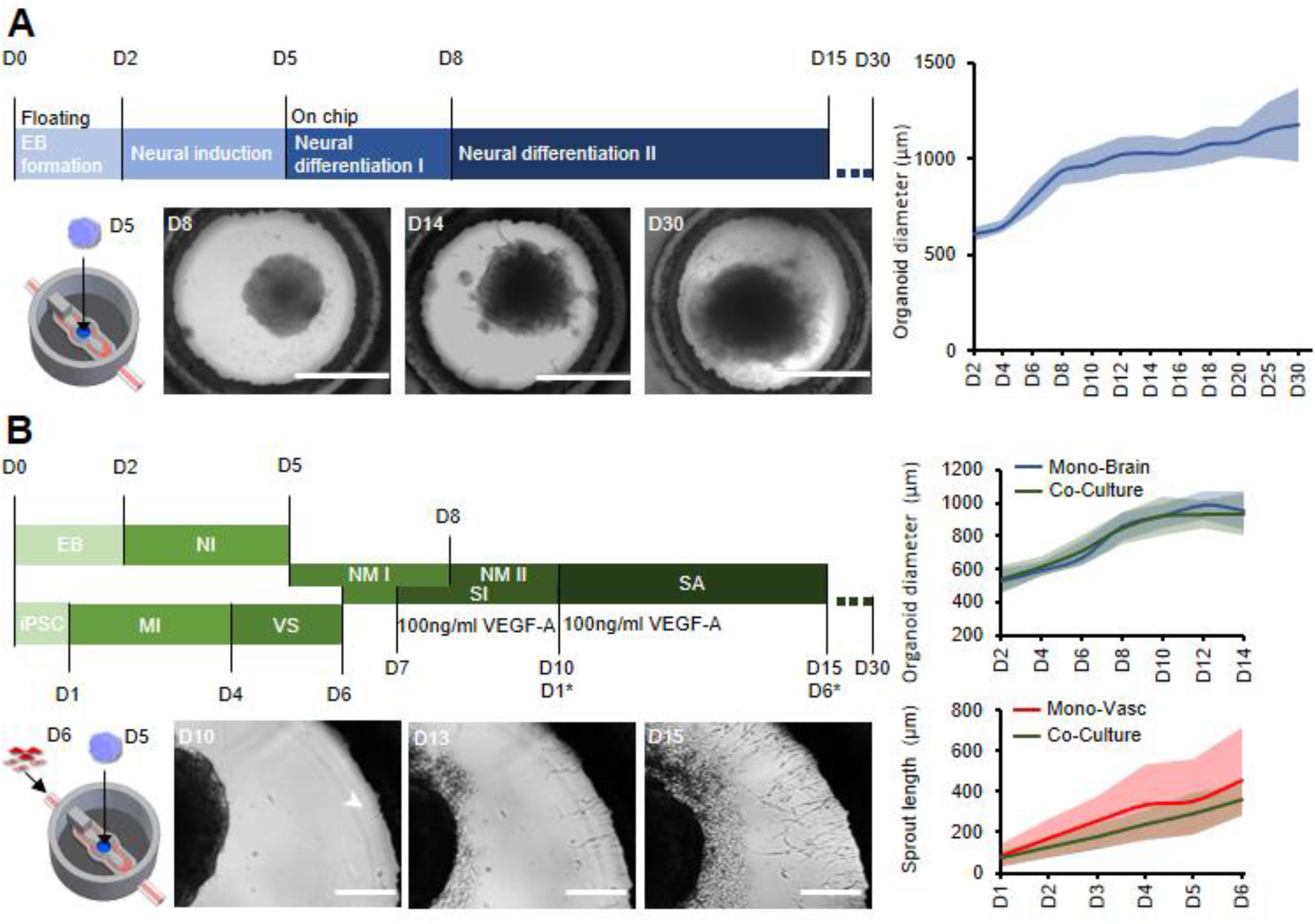
Neurovascular co-culture on chip. **(A)** Cerebral organoid differentiation protocol with brightfield images of Matrigel-embedded cerebral organoids on chip at days 8, 14 and 30 (scale: 1000μm) and quantification of cerebral organoid growth on chip (n=3;Avg.;SD) with 4-6 organoids analysed per n. (**B**) Neurovascular differentiation protocol on chip with bright field images of co-cultures on chip (scale: 250μm) and quantification of organoid size and sprout length (n=3;Avg.;SD) with 4-6 organoids analysed per n. (EB: embryoid body, NI: Neural Induction, NM: Neural maturation, MI: mesoderm induction, VS: vascular specification, SI: sprout induction, SA: Sprout analysis).

### Co-culture on 3D printed microfluidic chip

Having established that our platform was compatible with both vascular and cerebral organoid cultures, we next assessed whether these two cultures could be carried out together on-chip. We therefore developed a differentiation synchronization scheme whereby both vascular differentiation as well as cerebral organoid differentiation would be initiated off-chip, and would subsequently be brought together on chip at an early developmental time point.

Our protocol consisted in seeding cerebral organoids at day 5 of differentiation into the organoid culture chamber, and the vascular cells at day 6 of differentiation into the microfluidic channels (**Fig.3B**). In this way, co-differentiation would occur sufficiently late to allow for individual germ layer specification to occur, but early enough that feedback such as brain-specific endothelial specification to occur. In these conditions, both cultures showed good survival, with characteristic organoid growth in the central compartment and vascular sprouting from the channels. Sprout length was similar between mono and co-culture systems, indicating that the interaction between vascular cells and organoids did not alter sprouting characteristics. Conversely, cerebral organoid size in co-cultures was similar to that of organoids grown without vasculature, suggesting that the additional of vasculature did not impede organoid growth.

### Neurovascular organoid on 3D printed microfluidic chip

To characterize the molecular identity and localization of both neural and vascular cells in the initial co-culture period, the neurovascular constructs were stained at day 15 of differentiation, i.e. after 9 days of on-chip co-culture **(Fig.4A)**. Biii-tubulin+ post-mitotic neurons, CD31+ ECs, as well as PDGFB+ pericytes were all present in the co-culture at day 15, demonstrating that co-culture was permissive to the co-differentiation of these key cell types. Sprouts could be individually distinguished without overlap between each other, and were directed towards the organoid. To understand the three-dimensional tissue organization within the chip, we analysed individual slices of a 3D confocal image stack. ECs were seen to localize near the bottom of the microfluidic chip whereas pericytes and neurons were found approximately 40μm above the bottom of the plate (**Fig.4B**). Pericytes were observed in closer contact with the organoid, in contrast to ECs, suggesting that these cells rapidly migrated from the microfluidic channels or differentiated within the organoid and subsequently migrated out of the organoid.

**Figure 4.**
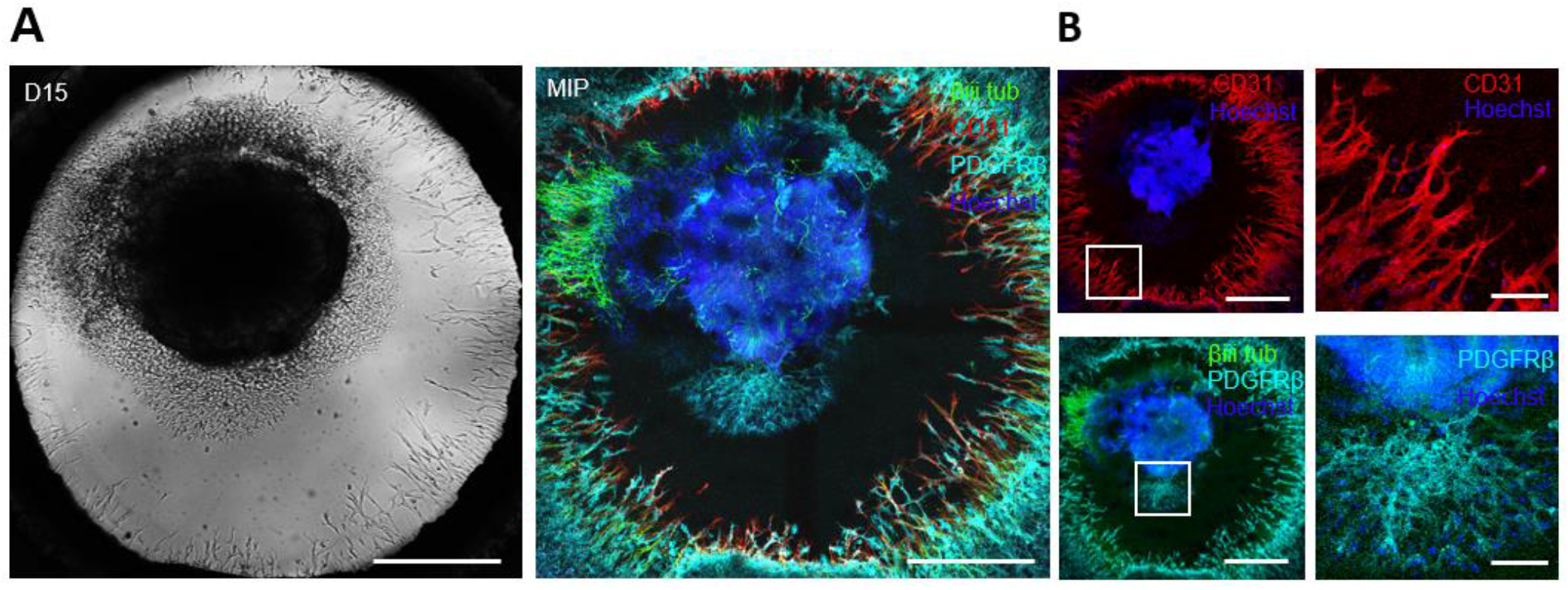
Neurovascular co culture on chip. (**A**) Brightfield image with corresponding 50μm MIP stained for βiii Tubulin, CD31 and PDGFRB of neurovascular culture at day 15 of differentiation (scale: 500μm). (**B**) Confocal sections of the MIP with CD31+ cells located at the bottom and βiii tubulin+ and PDGFRβ+ cells at 35 μm from the bottom slice (scale: 500μm, inset scale: 100μm), (MIP: Maximal intensity projection).

In order to verify that our platform could sustain longer-term interaction between organoids and vasculature, we maintained the co-cultures for up to 30 days of differentiation. Significant phenotypic changes were observed in the latter period of differentiation, including continued organoid growth concomitant with significant expansion and consolidation of the vascular network. In particular, vascular cells were observed to reach, and then surround, the central body of organoid. In multiple locations, endothelial cells were seen to invade the organoid body, as indicated by the co-localization of CD31+ cells with the dense nuclear signal from the core of the organoid. Strikingly, Biii tubulin+ neurites emanating from the central body of the neurovascular organoid frequently aligned with the vascular network. (**Fig5.A**). A three-dimensional reconstruction of the confocal images revealed that neural axons not only align with endothelial networks but also descend down from the organoid in order to come in contact with the vascular networks at the bottom of the microfluidic device (**Fig.5B**). This three-dimensional visualization additionally showed that these vascular networks formed an adjacent layer with the PDGFRβ+ pericytes (**Fig.5C**), suggesting the establishment of a complex patterned architecture linking pericytes, endothelial cells and cerebral organoid.

**Figure 5.**
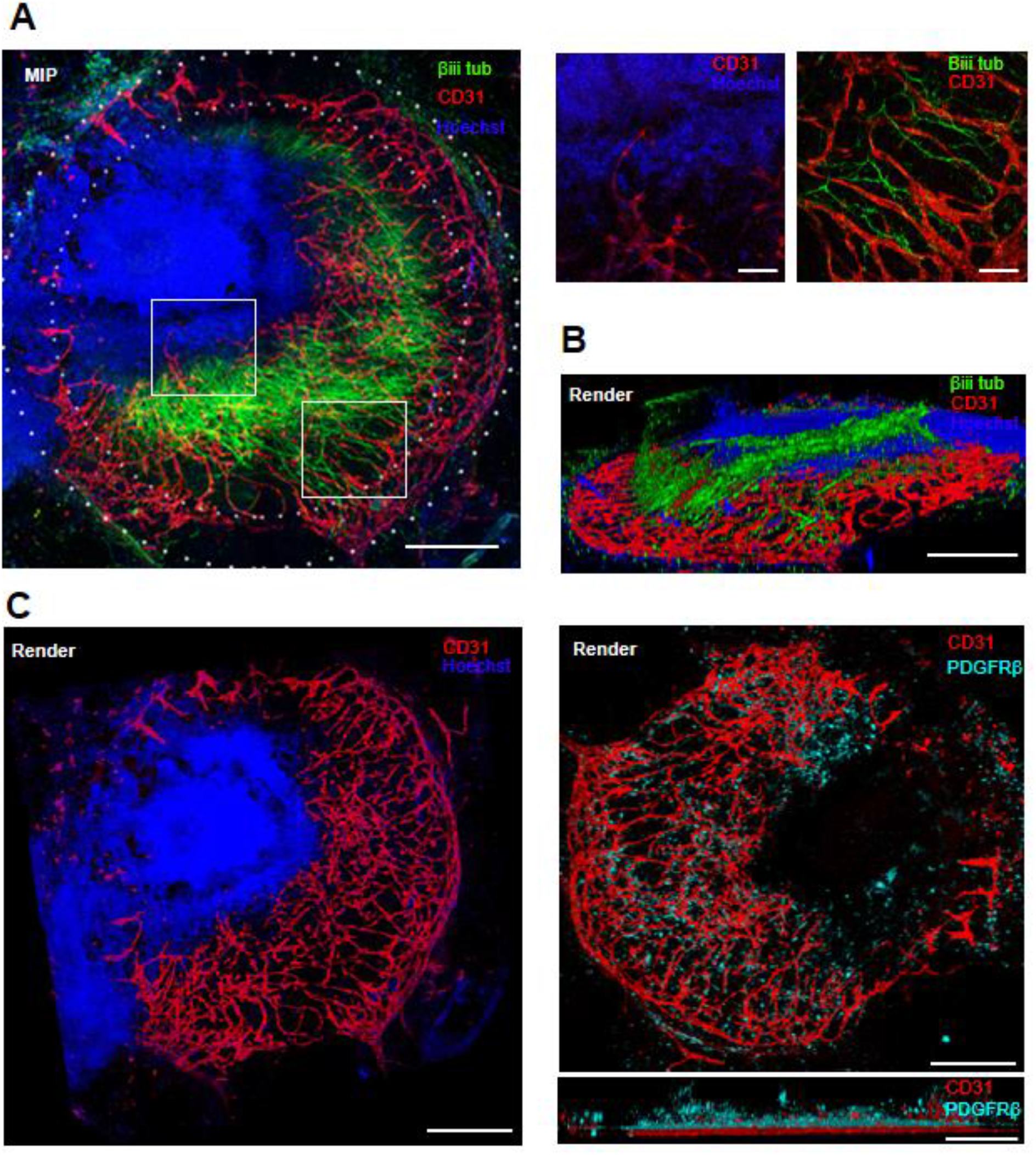
Neurovascular organoid on 3D printed microfluidic chip. (**A**) MIP of a 400μm confocal stack from a day 30 neurovascular organoid on chip IHC stained for βiii tubulin, CD31 and PDGFRB (scale: 500μm, inset scale: 100μm) (**B**) 3D render of 400μm confocal stack stained for βiii tubulin and CD31 (scale: 500μm). (**C**) 3D render of a day 30 neurovascular organoid on chip with CD31 vascular networks and PDGFRβ cells (scale 500μm). (MIP: Maximal intensity projection)

## Discussion

Here, we have demonstrated for the first time the possibility to control the interaction of endothelial cells, pericytes and organoids though the use of hPSCs and microfluidic technologies, in order to achieve an organoid vascularization process which is highly synchronized in time and space. The angiogenic sprouting and network formation observed in this platform are similar in size and branching architecture to those which have been reported with primary pericytes and ECs (11,35–38). An intriguing difference, however, is that primary cultures have been characterized by gel invasion of ECs first, followed by pericytes (38), whereas in our hPSC-based system simultaneous sprouting of ECs and pericytes is observed, reflecting possible PSC-specific characteristics or differences in developmental timing. We have also demonstrated that cerebral organoids growth and maturation within our chip, including the presence of characteristic Pax6+ and Biii-tubulin+ cells, which bear a close resemblance to previous studies where cerebral organoids were cultured both on a stationary PDMS micropillar plate and a PDMS microfluidic chip under flow condition (26,39). Our platform therefore demonstrates that organoids and vasculature can be grown in 3D printed chips, thereby opening up the organoid and vascular biology fields to the possibilities offered by the design flexibility of 3D printing. Indeed, the significant price drops in recent years in both hardware and materials of such 3D printing technologies have made them readily accessible to every lab (25).

The generation of on-chip interactions between hPSC-derived organoid and vasculature leading to a complex neurovascular unit demonstrates the possibilities of this platform to more precisely investigate the role of ECs and pericytes in the development of human tissue. Our standardized on-chip vascularization approach overcomes a number of limitations with other recently reported technologies (40). For example, subtractive removal of material from organoid tissue by sacrificial laser writing is a promising technique for creating perfusable channels (41), however the process of tissue ablation also has clear implications in altering tissue development. The use of a manufactured chip-like co-culture system on a re-engineered 384 well plate to study organoid-vasculature has also been recently proposed in the context of interaction between colon organoids and HUVECs and fibroblasts (11). This parallelized approach suggests that this technique could be useful for high throughput screening of drug candidates, however the use of such a standardized plate limits users flexibility in terms of the spatial control, which can be overcome using accessible 3D printing technology.

The organoid vascularization approach we have shown here is designed to be generic and widely applicable, and opens a number of avenues for further research in the field of in-vitro tissue vascularization. We expect that our protocol for generating ECs and pericytes from hPSC, in combination with PSC-derived organoids systems, will allow for a broader use of such developmentally synchronized co-cultures. Additionally, on-chip vascularization where the vascular and organoid components are brought together at defined times and allowed to interact starting from a defined spatial relationship could be a widely applicable strategy to generate a variety of other vascularized organoids. More broadly, the design and 3D printing of customizable microfluidic devices with consumer-grade printers and appropriately chosen and processed biocompatible materials may pave the way towards wider access and use of such technologies to generate controllable and reproducible *in vitro* tissues.

## Acknowledgements

This work was supported by the FWO grant G087018N and FWO postdoctoral fellowship 1217220N, Interreg Biomat-on-Chip grant and Vlaams-Brabant and Flemish Government co-financing, KU Leuven grants C14/17/111 and C32/17/027 and King Baudouin Foundation grant J1810950-207421.

## Author Contributions

IS conducted experiments and analysis. SG,AA and GR conducted experiments. IS and TP developed the FDM 3D printed holders. IS, SG,AA, AR and CV interpreted the data and edited the manuscript. IS and AR wrote the manuscript.

## Supplementary Figures

**Supplementary Figure 1.**
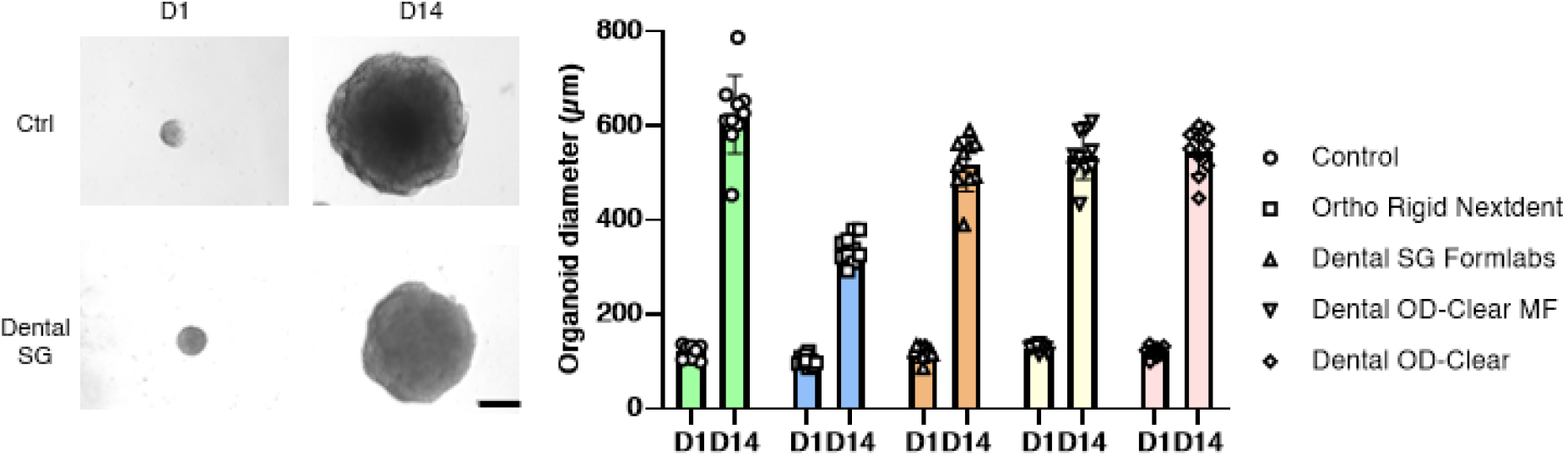
Organoid growth on cured resins. Brightfield images of organoids at D1 and D14 on top of cured resin (scale: 200μm) and organoid diameter quantification with 10 organoids measured per condition.

**Supplementary Figure 2.**
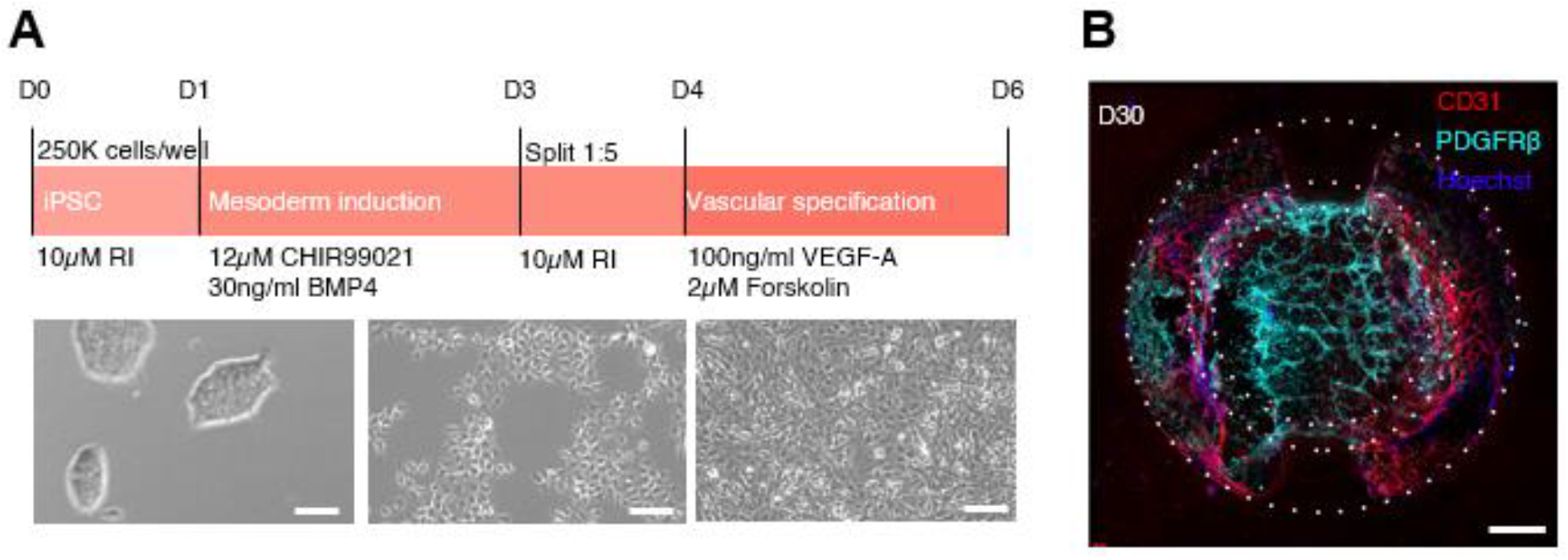
Directed differentiation of hIPSC into pericytes and endothelial cells. (**A**) Vascular differentiation protocol with brightfield images taken at D2,D4 and D6 during 2D vascular differentiation of hIPSC (scale: 100μm). **(B)** IHC of D30 organoid 75μm from coverslip (scale: 500μm).

**Supplementary Figure 3.**
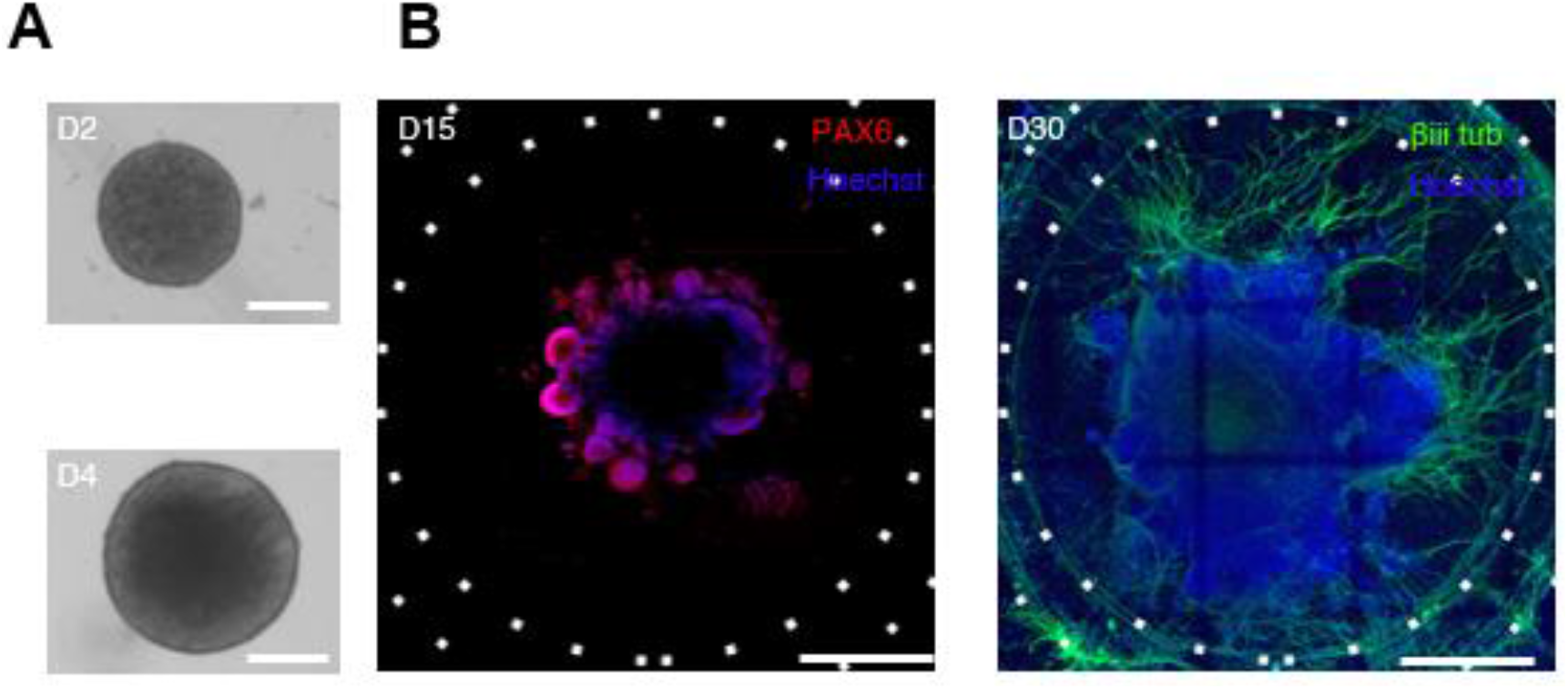
Directed differentiation of hIPSC into cerebral organoids on 3D printed microfluidic chips. (**A**) Brightfield images of aggregated cells at two days after aggregation and neural aggregate four days after aggregation (scale: 250μm), and (**B**) IHC of PAX6 and βiii tubulin of D15 and D30 Cerebral organoids on chip (scale: 500μm).

